# Anti-amyloid antibody effects on Aβ-42 protein aggregates profiled using nanospectroscopy

**DOI:** 10.1101/2025.07.17.665305

**Authors:** Nico Kummer, Martina Cihova, Peter Niraj Nirmalraj

## Abstract

Anti-amyloid beta (Aβ) drugs such as aducanumab and lecanemab are designed to clear brain amyloids and slow Alzheimer’s disease (AD) progression. While immunoassays provide ensemble-level details on anti-Aβ drug interactions with protein targets, their interfacial effects are largely unknown at a single-particle level. Here, we profile untreated and aducanumab-treated Aβ-42 protein aggregates from oligomers to fibrils using atomic force microscopy coupled with infrared spectroscopy (nanospectroscopy). Based on the recorded morphological and secondary structure details of aducanumab-treated Aβ-42 aggregates using nanospectroscopy, we observed a reduction in oligomer prevalence and formation of larger diameter fibril bundles compared to identically prepared untreated-Aβ-42 peptides. Conversely, controls based on lecanemab did not reveal any quenching of the Aβ-42 oligomer generation. Moreover, lecanemab was evidenced to bind along the full length of the Aβ-42 protofibril surface preferentially. Importantly, the structure of Aβ-42 fibrils did not disassemble in both studies upon the adsorption of aducanumab and lecanemab. Additional experiments were also conducted, such as aggregation kinetic assays and Fourier transform infrared spectroscopy on aducanumab and lecanemab-treated Aβ-42 protein aggregates that corroborated the findings from nanospectroscopy studies. Our work highlights the usefulness of nanospectroscopy in studying elemental anti-amyloid antibody interactions with protein biomarkers, a requisite for improving Alzheimer’s disease-modifying treatments.

## Introduction

Alzheimer’s disease (AD) is the most prevalent form of dementia in aging societies^1^ and is a progressive neurodegenerative disorder characterized by declining memory and cognitive abilities in individuals. While AD accounts for about 70% of dementia cases, it rarely occurs in isolation, with most patients exhibiting mixed pathologies, such as Parkinson’s disease, frontotemporal, and vascular dementia. AD is histopathologically evidenced by the accumulation of amyloid beta (Aβ) and tau protein deposits in brain tissue. The molecular pathogenesis of AD is known to onset nearly 20 years before symptoms such as loss of memory and cognition become visible^2^. Although AD is highly complex and is prone to multifactorial triggers^3^ spanning from aging, genetic to environmental risk, the degree of aggregation of specific amyloid and tau isoforms can reflect disease severity and hence can be used as biomarkers to monitor disease progression^4^. In particular, measuring the ratio of Aβ-40 to Aβ-42 as well as total-tau (t-tau) and phosphorylated-tau (p-tau) levels in cerebrospinal fluid (CSF) and plasma using biochemical assays has been demonstrated to be a potential indicator of cerebral amyloid and tau pathology in AD dementia^5^.

Currently, there are no known therapeutic interventions to cure AD or reverse cognitive decline in patients. However, the American Food and Drug Administration (FDA) has previously approved monoclonal anti-amyloid antibodies such as aducanumab^6^, lecanemab^7^, and donanemab^8^ based on evidence that these compounds can recognize and clear protein aggregates through phagocytosis by microglia in brain tissue^9^. Aducanumab (human IgG1 anti-Aβ monoclonal) was one of the first in the series of anti-amyloid antibodies to specifically target both soluble and insoluble forms of amyloids and reduce brain amyloid aggregates confirmed through structural brain imaging ^10^. However, the manufacture of aducanumab has been discontinued by Biogen, citing reprioritizing resources to advance efforts related to lecanemab. Conversely, research aimed at understanding the elemental effects of aducanumab on amyloid aggregates continues, which is crucial for the future design of therapeutic agents for treating and managing AD^11^. In this context, phase III trials on another anti-amyloid antibody, Lecanemab, have been shown to target specific polymorphs of Aβ and to limit cognitive decline in patients with mild cognitive impairment and early AD symptoms. It is known that aducanumab and lecanemab both have an affinity to bind to the higher-order aggregates (i.e., oligomers, protofibrils, fibrils, and plaques) of Aβ rather than the monomers^12^. While aducanumab showed a specific affinity to mature fibrillar aggregates, lecanemab moderately binds to plaques^12^ but has increased preferential binding to protofibrils^12^, which is an intermediate state characterized by a nodular morphology that occurs before the onset and formation of mature fibrils^13, 14^. Studies based on kinetic analysis combined with quantitative binding measurements have also shown that aducanumab binds to the surface of Aβ-42 and reduces the secondary generation of related oligomers^15^. In parallel, studies using immunoassays and molecular dynamics simulations have elucidated the binding mechanism of aducanumab to amino acid residues 3-7 of Aβ through interactions between a hydrophobic pocket and the epitope^16, 17^. In addition to understanding AD progression and quantifying the effect of therapeutic agents on clinically relevant protein biomarkers using ensemble-based techniques such as chemiluminescent-based immunoassays and fluorescence-based binding assays to map the protein aggregation kinetics in a high-throughput manner, label-free and single-particle level detection techniques are also fast gaining traction to provide new insights on AD fluid biomarkers.

To this end, atomic force microscopy (AFM), has been demonstrated to be a powerful imaging tool in resolving and quantifying the full spectrum of aggregates formed along the primary^14, 18, 19^ (fibril-independent, and secondary nucleation^20^ (fibril-dependent) pathway of Aβ-40, Aβ-42 and tau proteins under standard laboratory conditions at the single-particle level. Furthermore, the interfacial interactions of lecanemab with surface-confined Aβ aggregates have also been resolved in real-time using video-rate AFM^21^. There is also an emerging trend for employing AFM operating under standard laboratory conditions to directly resolve protein aggregates in body fluids and correlate the measured aggregate morphology with neurological disease progression^22-24^. In the context of studies related to AD it has been demonstrated that soluble oligomers extracted from AD brain tissue can be characterised using AFM, and distinctions were shown between spherical and elongated aggregates, which is crucial information in tracking disease progression and potentially identifying therapeutic targets^25^. In our laboratory, we have used AFM operating in a liquid environment to resolve protein aggregates on the surface of red blood cells^26, 27^ and in the CSF^13^ of AD patients. From our previous work, we have observed the prevalence of fibrillar aggregates and fibril length to correlate with the various stages of AD^13, 26^. AFM not only provides information on the morphological information but also allows measuring other properties such as stiffness, surface potential, and even information on secondary structure of protein aggregates in combination with localized chemical spectroscopy. In particular, infrared (IR) spectroscopy is a well-established technique capable of providing chemical information on the secondary structure of protein aggregates. Taken together, the combination of AFM with IR spectroscopy is a robust methodology to jointly study the morphology and chemical structure of proteins implicated in AD. Such an approach relies on the photothermal effect, where infrared laser radiation is absorbed in a sample, converted into heat, and subsequently leads to a local thermal expansion, which can be detected using an AFM cantilever^28-31^. Such a nanoimaging and spectroscopy approach has been previously demonstrated to obtain chemical maps at selected wavenumbers in the amide I region and localized infrared spectra on individual amyloid nanofibrils of the aggregated Josephin domain of ataxin-3 implicated in spinocerebellar ataxia type 3^32^. More recently, AFM-IR has also been extended to characterize Aβ^33-42^, tau^43^, and α-synuclein^34, 37, 42^ proteins. The findings from nanospectroscopy^40^ were in good agreement with previous FTIR spectroscopy results^44^ referring to anti-parallel beta-sheet Aβ-42 oligomers as the origin of parallel beta-sheet fibrillar aggregates at fast kinetic rates. Furthermore, the inhibitory effect of the small molecule bexarotene on amyloid-β aggregation has been elucidated with AFM-IR^45^ and complementary scattering scanning near field optical microscopy (s-SNOM)^46^. AFM-IR has also been used to resolve Aβ and α-synuclein aggregates extracted from human brain tissue^42^, highlighting the versatility of this nanoscale analytics approach.

In this paper, we resolve in a label-free manner the differential effects of aducanumab and lecanemab (anti-Aβ antibodies) on synthetically prepared Aβ-42 protein aggregates using AFM-IR at a single-particle level. AFM-IR measurements provide information about protein secondary structure, which we demonstrate is a useful parameter to compare anti-Aβ antibody-treated and untreated Aβ-42 protein aggregates. While untreated Aβ-42 fibrils predominantly consist of parallel β-sheets, the addition of anti-Aβ antibodies at the beginning of the aggregation leads to a measurable increase in α-helices. The findings at the nanoscale agree well with bulk FTIR spectra showing the same shift in and secondary structure. In general, the nanospectroscopy-based imaging approach confirms that, compared to lecanemab, only aducanumab tends to significantly reduce the production of free Aβ-42 oligomers. Furthermore, lecanemab was observed to have a specific binding affinity along the entire length of the Aβ-42 protofibrils, which is useful information for designing drugs for targeted therapeutics.

## Results and Discussion

First, we studied the diverse morphologies of Aβ-42 protein aggregates incubated for 24 h at 37°C (see materials and methods section for details on peptide concentration and AFM sample preparation) using standard AFM. Figure 1A is an AFM topographic image showing the presence of fibrillar and spherical Aβ-42 particles on a gold surface. To calculate the diameter of the single particles and elongated cylindrical fibrils, cross-sectional profiles were extracted across the particles as indicated in the AFM image (Fig. 1A). Figure 1B shows a set of color-coded cross-sectional profiles measured along single fibrils marked by red, green, and black lines in Fig. 1A. Height profiles extracted across spherical particles (marked by red and black circle in Fig.1A), which could potentially represent the oligomers is shown in Fig. 1C. Since the measured width of a particle represents the convolution of the fibril topography and the tip geometry, the extracted height difference between the surrounding substrate and the highest point on a spherical and cylindrical particle is treated as the particle diameter. Additional information is provided in Fig. S1, supporting information for height profiles obtained from Aβ-42 spherical particles. In contrast to pure Aβ-42 aggregate morphology, incubating identically prepared Aβ-42 peptides in the presence of aducanumab led to the aggregation of individual fibrils to form bundles with a larger height compared to single fibril and were observed to have fewer spherical particles, as shown from a representative high-resolution AFM topograph (Figure 1D). The surface profile of fibril bundles appears corrugated due to adsorbed aducanumab antibodies and were measured to have diameter of ∼20 nm as seen in the extracted cross-sections (Figure 1E), whereas the diameter of the spherical particles did not vary significantly compared to spherical particles previously resolved and quantified from the AFM topographs for the Aβ-42 untreated samples (Figure 1F). To qualitatively assess if aducanumab binds to the hydrophobic patches in the primary sequence of Aβ-42 (cryo-electron microscopy-based atomic structure, PDB code: 5OQV, Figure 1G), surface potential measurements on individual untreated and aducanumab-treated Aβ-42 fibrils were performed using Kelvin probe force microscopy (see materials and methods section for details on KPFM and imaging protocols and Figure 1H-K and supporting information Figure S2-4 for details on KPFM control measurements). Previously, surface potential measurements using KPFM have shown typical surface potential dependence of Aβ on pH and allowed the identification of the isoelectric point^47^. KPFM was used here for its ability to provide spatially resolved information of surface potential at the nanometer scale, making it a powerful tool for probing electrostatic heterogeneity along individual fibrils ^47, 48^. The untreated Aβ-42 samples showed a relatively smoother fibril surface (Figure 1H) and only moderate changes in surface potential along the fibril length (Figure 1I).

**Figure 1:**
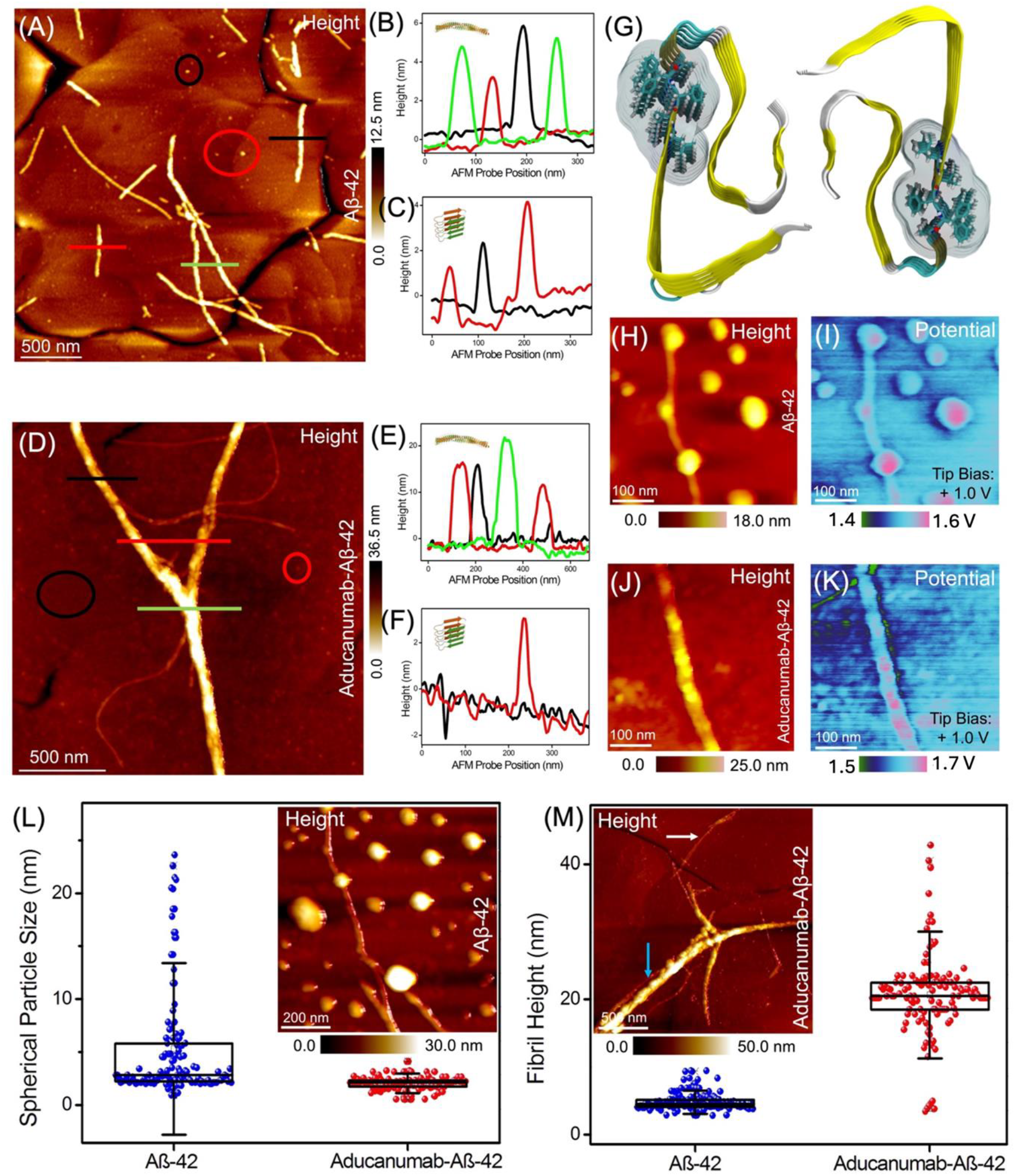
Morphological characterization of aducanumab-treated and untreated Aβ-42 protein aggregates. (A) AFM height image of untreated Aβ-42 incubated for 24 h deposited on a gold substrate. (B) Representative height profiles of fibrillar aggregates extracted along the black, red, and green lines in (A). (C) Representative height profiles of oligomeric particles in the areas encircled in black and red in (A). (D) AFM height image of aducanumab treated Aβ-42 incubated for 24 h deposited on a gold substrate. (E) Representative height profiles of fibrillar aggregates extracted along the black, red, and green lines in (D). (F) Representative height profiles of oligomeric particles in the areas encircled in black and red in (D). (G) Cryo-electron microscopy resolved the atomic structure of Aβ-42 fibril (PDB identifier code: 5OQV^55^). The shaded area represents the hydrophobic patches along the fibril backbone. AFM height image of a single untreated Aβ-42 fibril incubated for 4 h deposited on a gold substrate. KPFM surface potential map corresponding to (H). (J) AFM height image of a single aducanumab treated Aβ-42 fibril incubated for 4 h deposited on a gold substrate. (K) KPFM surface potential map corresponding to (H). (L) Box diagram showing the mean spherical particle size of treated and untreated Aβ-42 after 24 hours of incubation including standard deviation and 25% and 75% percentiles, the inserted AFM height image shows abundant spherical particles. (M) Box diagram showing the mean fibril height of treated and untreated Aβ-42 including standard deviation and 25% and 75% percentiles, the inserted AFM height image shows a fibril bundle (blue arrow) and a single fibril (white arrow) found in Aβ-42 samples treated with aducanumab.

However, differences in surface potential between the fibril surface and the adsorbed spherical particle are evident from the surface potential map (Fig. 1I), with potential variations on the order of 20-50 mV (Fig. S4, Supporting Information). In the treated samples, the fibril surface had a higher roughness due to the adsorbed aducanumab particles (Figure 1J). Although the surface potential profile exhibited more distinct spatial variation along the fibril length, the magnitude of intra-fibril potential fluctuation (∼30 mV, see Supporting Information Fig. S4) did not differ significantly from that observed in the untreated sample (Figure 1K, Fig. S4, Supporting Information). The surface potential fluctuations observed across both untreated and aducanumab-treated Aβ-42 fibrils on the order of low-to-mid tens of millivolts are consistent with earlier KPFM studies on amyloid assemblies, including fibrillar Aβ-42 aggregates at physiological pH^47, 48^. Similarly, the zeta potential measured for Aβ-42 fibrils in PBS medium was ™19.2 mV and was only reduced slightly to ™14.6 mV after aducanumab treatment. These results suggest that charge played a negligible role and that the binding mechanism was potentially based on hydrogen bonding and hydrophobic interactions, as previously reported in the literature^16, 17^. In particular, it was demonstrated, using ThT fluorescence kinetic assays, that aducanumab tends to bind to hydrophobic patches of Aβ-42 fibrils, reduces the rate of the secondary nucleation pathway, and reduces free oligomer concentration^49^. Based on the AFM measurements on untreated and aducanumab-treated Aβ-42 protein aggregates, we quantified the complete spectrum of aggregate morphologies. Figure 1L is a statistical plot showing the differences in spherical particle size for untreated (coded in blue) and aducanumab-treated Aβ-42 (coded in red). The quantified AFM data indicate that aducanumab-treated Aβ-42 aggregates contained spherical particles of a lower diameter compared to untreated-Aβ-42 protein aggregates. The formation of Aβ-42 fibril bundles after aducanumab treatment is also confirmed by the measured higher mean fibril height (Figure 1M). The inserted AFM height image in Fig. 1M clearly shows the differences between single fibrils (marked by a white arrow) and thicker fibril bundles (marked by a blue arrow) within a fixed imaging frame area. The heterogeneity in the surface structure of the fibrils and the differences in the packing of fibrils within the bundles, resolved using AFM, could also have a minor contribution to the broad distribution of the measured fibril heights, as shown in Fig. 1M. See Fig. S5 in the supporting information shows high-resolution AFM images of aducanumab-treated Aβ-42 fibrils. Standard AFM measurements provide morphological details, and KPFM-based analysis provides insights into differences in surface potential between fibrillar and non-fibrillar components. These imaging approaches have limitations. For instance, it is not feasible to unambiguously distinguish between single oligomers and aducanumab particles adsorbed on the fibril surface based solely on AFM height profiles, and KPFM conducted under ambient conditions could be subject to environmental artifacts^50^.

To address these experimental challenges, we conducted AFM-IR measurements (see materials and methods section for details on AFM-IR tool and imaging protocols), which can simultaneously obtain morphological and chemical structure information under standard laboratory conditions. First, we analysed purified Aβ-42 aggregates using AFM-IR on samples that were drop-casted and air-dried on an ultra-flat gold surface to gain information about their chemical structure. AFM-IR uses a tunable infrared laser focused on the Aβ-42 aggregates adsorbed on a gold substrate under the AFM tip. (Figure 2). When the wavenumber of the laser matches an absorption band of the sample, the absorbed radiation is proportional to the thermal expansion caused by the photothermal effect. Keeping the wavenumber constant at an absorption band of interest and scanning over a sample area allows chemical mapping. By sweeping through a wavenumber range, the response of the AFM probe can be converted into an IR absorption spectrum, giving chemical information about protein aggregates. See supporting information Figs S6-7 for details on AFM-IR data analysis protocols used in the treatment of the IR spectra. Figure 3A shows a tapping mode AFM-IR height image with fibrillar and spherical Aβ-42 aggregates, which confirms the previously observed findings using a standard AFM (Figs.1A-C). The phase image (Figure 3B) and the IR map at 1630 cm^-1^, a wavenumber within the amide I region (1600-1700 cm^-1^) corresponding to the signal from parallel β-sheets (Figure 3C) are overlaid in Figure 3D to demonstrate the good correlation of the IR signal with the pure AFM measurement channel. The fibrillar aggregates show a strong IR signal, and there is also a detectable signal from the smaller spherical particles present in the background. Conversely, the larger spherical particles resolved in the AFM topograph do not give any IR signal, suggesting that these particles are not proteinaceous. Instead, they could be salt residues that were not rinsed off during sample preparation. It is known from previous studies that the AFM-IR tool, when operated in contact mode, results in a better signal-to-noise ratio, especially when collecting reproducible spectra^40, 45^ from single amyloid fibrils. To verify if there is a difference in the quality of the recorded IR spectra obtained in tapping mode, we repeated the AFM-IR experiments in contact mode. Figure 3E shows a contact mode AFM-IR height image with fibrillar Aβ-42 aggregates and corresponding IR maps at 1636 cm^-1^, 1668 cm^-1^, and 1695 cm^-1^, respectively (Figure 3 F-H). The IR signal was the most intense for the map at 1636 cm^-1^, showing the contribution from parallel β-sheets, followed by a lower intensity at 1668 cm^-1^, corresponding to α-helix contribution; the signal from anti-parallel β-sheets (1695 cm^-1^) was not detected. This is in good agreement with the corresponding spectral data shown in Figure 3I, which shows a pronounced peak at 1630 cm^-1^ within the amide I region. The spectra have been de-spiked and treated with Savitzky-Golay smoothing; the raw, de-spiked, and final smoothed spectra are shown in Supplementary Information, Figure S6. The second derivative of the amide I peak is dominated by the minimum at around 1630 cm^-1^ (Figure 3F), corresponding to the contribution of parallel β-sheets. The weak minima around 1660 cm^-1^ show a negligible contribution from α-helical secondary structure. FTIR experiments show a sharp peak around 1630 cm^-1^ (supporting information, Figure S7) and are in good agreement with the literature^38, 40^, thereby confirming the presence of mostly parallel β-sheets.

**Figure 2:**
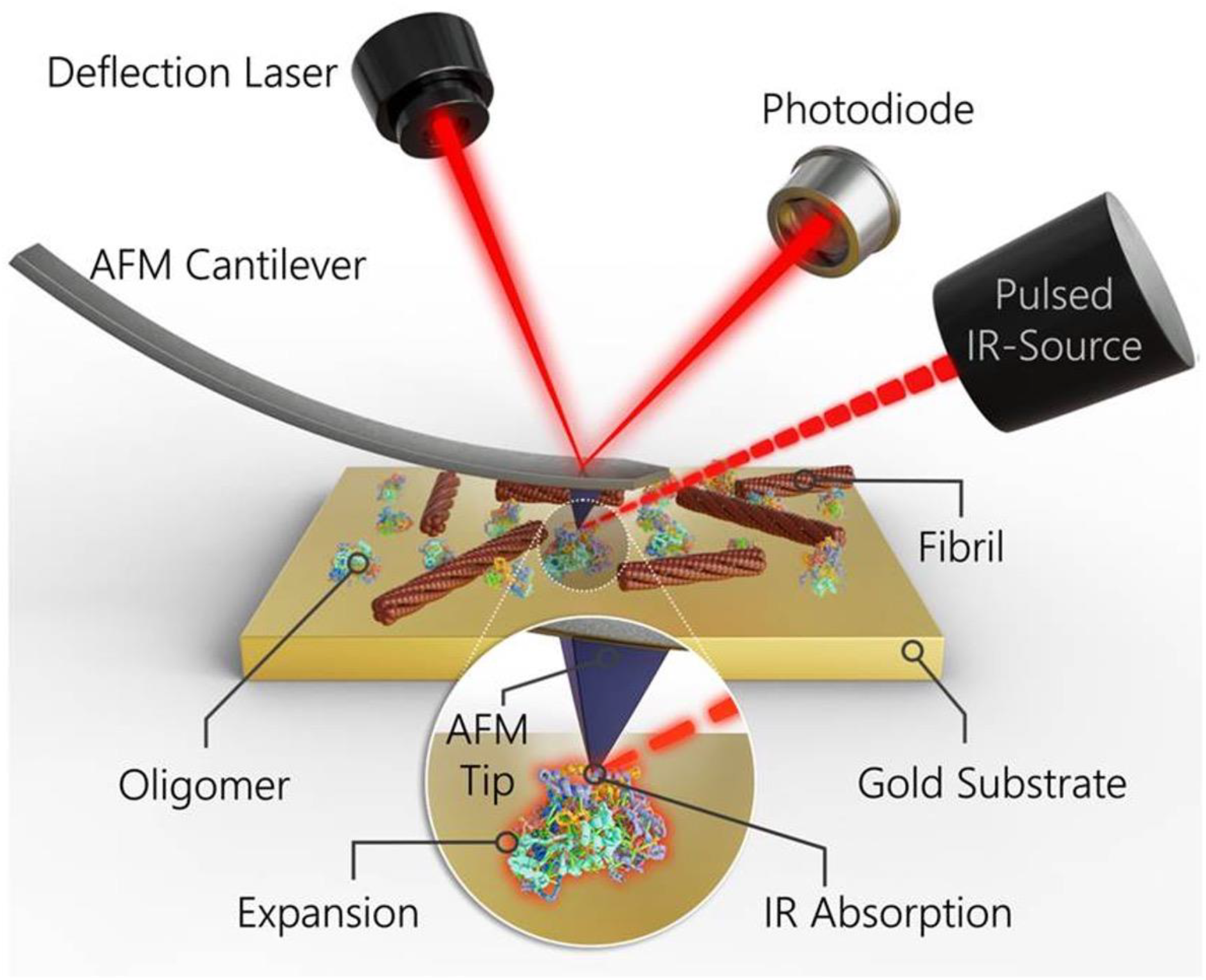
Working principle of AFM-IR measurements on Aβ-42 aggregates. The pulsed IR laser is directed at the sample, in which the absorption of IR light leads to photothermal heat generation and expansion of the Aβ-42 adsorbed on a gold substrate. The expansion can be measured with the AFM system: the deflection of the cantilever is measured by a photodiode detecting light from the deflection laser reflected on the back of the cantilever (objects are shown not to scale).

**Figure 3:**
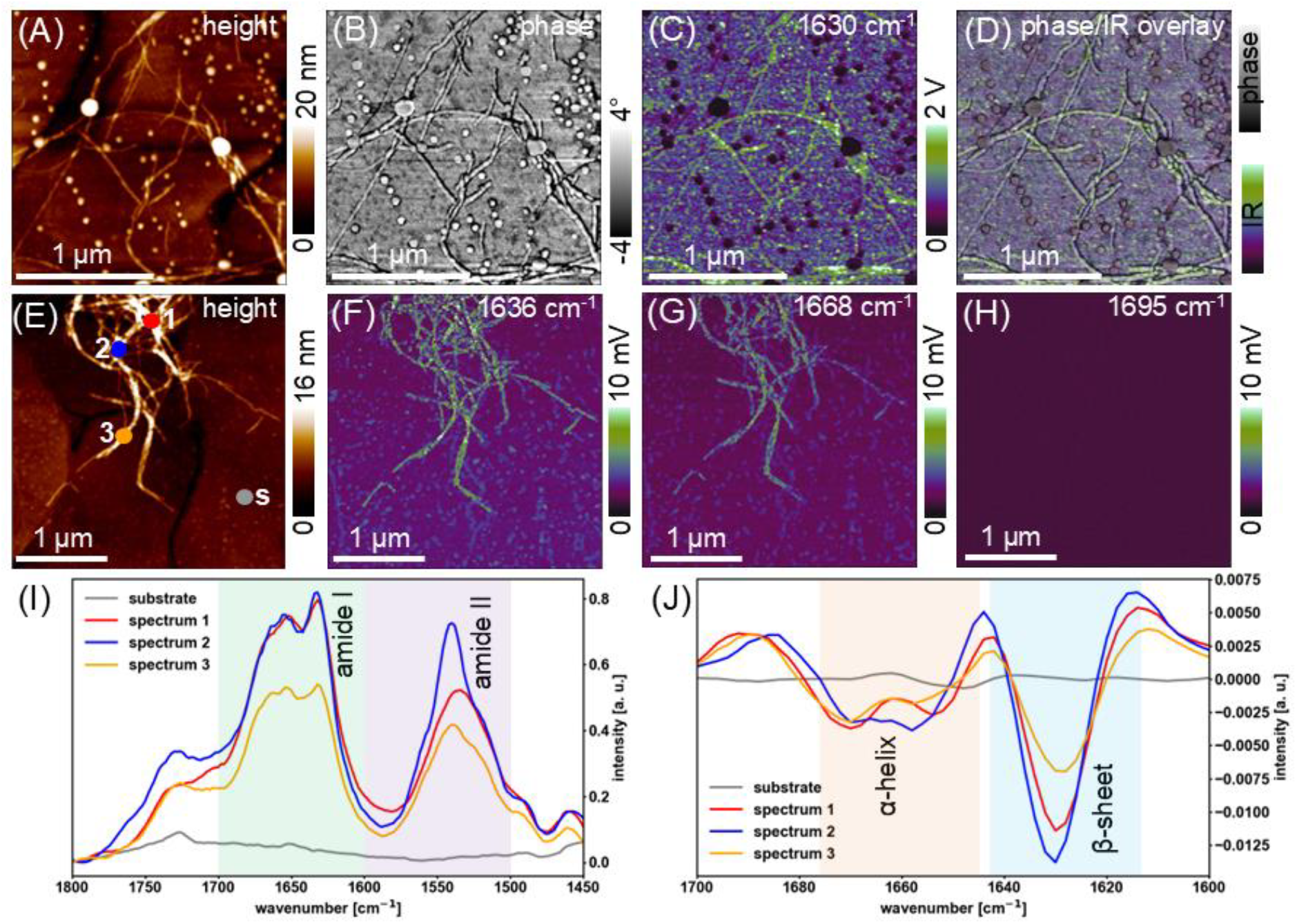
Nanospectroscopic analysis of Aβ-42 protein aggregates using tapping and contact mode AFM-IR. (A) Tapping mode AFM-IR height image of untreated Aβ-42 incubated for 24 h, deposited on a gold substrate. (B) Corresponding tapping mode phase image. (C) Corresponding IR map at 1630 cm^-1^, a wavenumber within the amide I region showing the contribution from parallel β-sheets. (D) Overlay of the phase image and IR map. (E) Contact mode AFM-IR height image of untreated Aβ-42 incubated for 24 h, deposited on a gold substrate. (F-H) Corresponding IR maps at 1636 cm^-1^, 1668 cm^-1^, and 1695 cm^-1^, wavenumbers within the amide I region showing the contribution from parallel β-sheet, α-helix, and anti-parallel β-sheet secondary structure, respectively. (I) Representative local AFM-IR spectra for the substrate (marked “s” in the height image) and three fibrils (marked “1-3”) showing the amide I and II regions (1450-1800 cm). (J) Second derivatives of the spectra showing the amide I region, the minima correspond to the contribution from α-helix and β-sheet secondary structure.

Upon developing a reliable and reproducible AFM-IR imaging protocol using purified Aβ-42 protein aggregates, we asked the question of whether secondary structural changes can be detected using our approach when Aβ-42 peptides are incubated together with aducanumab. Figure 4A is an AFM height image of aducanumab-treated Aβ-42 fibrillar aggregates recorded in tapping mode, and the corresponding phase signal is shown in Fig. 4B. Both the height and phase maps confirm the presence of aggregated fibrils. Figure 4C shows the corresponding IR map at 1636 cm^-1^ (parallel β-sheet contribution) with a strong signal from the bundles and a weaker signal from spherical particles in the background, which correlated well with the phase, as shown in the overlay (Figure 4D). In the contact mode measurements on aducanumab-treated Aβ-42 bundles (Figure 4E), very similar signal intensities were measured for parallel β-sheets at 1636 cm^-1^ (Figure 4F) and α-helices at 1668 cm^-1^ (Figure 4G), while the anti-parallel β-sheet contribution at 1695 cm^-1^ (Figure 4H) was weak. The IR maps were in good agreement with the amide I peak in the absorption spectra on the fibril bundles, as displayed in Figure 4I (raw spectra can be found in Supporting Information, Figure S8), which only had a weak shoulder instead of a pronounced peak at 1630 cm^-1^, hinting towards a lower parallel β-sheet content. This became more evident in the second derivatives of the amide I peak (Figure 4F), exhibiting similar minima for parallel β-sheets and α-helices, confirming the binding of aducanumab, which is rich in an α-helical structure. This is in good agreement with the lower β-sheet peak in the bulk FTIR spectrum of the aducanumab-treated Aβ-42 after 0.5 and 24 h, shown in Supporting Information, Figure S7A and B. The absence of a pronounced β-sheet peak in aducanumab-treated Aβ-42 after 0.5 hours, while the untreated sample already shows β-sheet formation, suggests that aducanumab inhibits aggregation. Next, measurements using thioflavin T fluorescence kinetics assays confirmed the lower rate of aggregation when Aβ-42 peptides were treated with aducanumab (see Supporting Information, Figure S9). Our findings based on AFM, KPFM, AFM-IR, and aggregation kinetic assays are in agreement with previous kinetic studies that aducanumab tends to slow down the formation of oligomers^49^.

**Figure 4:**
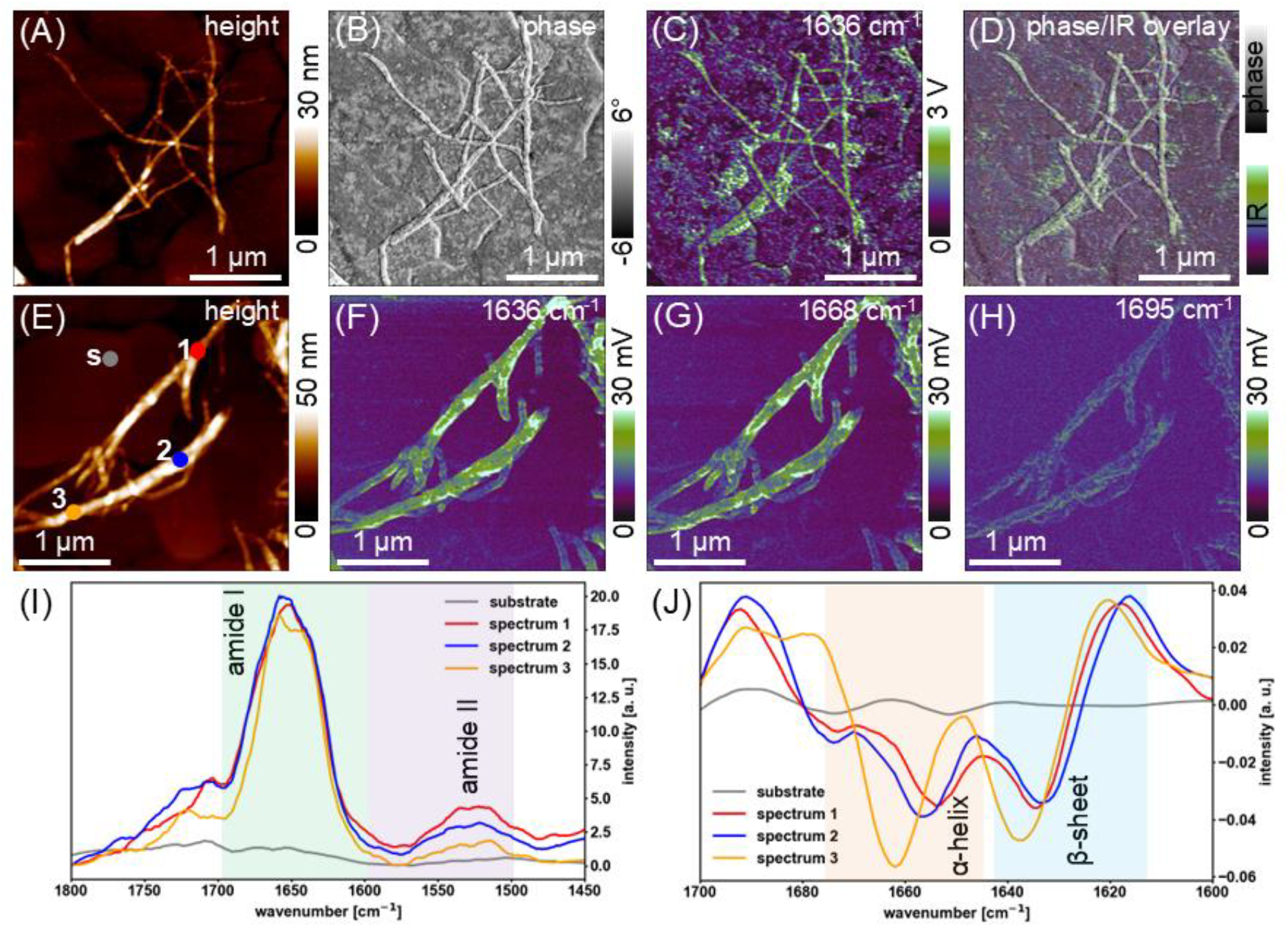
Nanospectroscopic analysis of Aβ-42 protein aggregates treated with aducanumab using tapping and contact mode AFM-IR. (A) Tapping mode AFM-IR height image of aducanumab treated Aβ-42 incubated for 24 h deposited on a gold substrate. (B) Corresponding tapping mode phase image. (C) Corresponding IR map at 1636 cm^-1^, a wavenumber within the amide I region showing the contribution from parallel β-sheets. (D) Overlay of the phase image and IR map. (E) Contact mode AFM-IR height image of aducanumab treated Aβ-42 incubated for 24 h deposited on a gold substrate. (F-H) Corresponding IR maps at 1636 cm^-1^, 1668 cm^-1^, and 1695 cm^-1^, wavenumbers within the amide I region showing the contribution from parallel β-sheet, α-helix, and anti-parallel β-sheet secondary structure, respectively. (I) Representative local AFM-IR spectra for the substrate (marked “s” in the height image) and three fibrils (marked “1-3”) showing the amide I and II regions (1450-1800 cm). (J) Second derivatives of the spectra showing the amide I region, the minima correspond to the contribution from α-helix and β-sheet secondary structure.

To compare the binding of different monoclonal antibodies, we repeated the AFM and AFM-IR measurements on Aβ-42 aggregated in the presence of lecanemab, which is currently a promising drug candidate in the treatment of AD^51^. Similar to aducanumab, the addition of lecanemab at the beginning of the aggregation led to agglomeration and the formation of fibril bundles, as shown in Figure 5A and B. Treatment of Aβ-42 with lecanemab also resulted in a corrugated fibril surface, as revealed in the high-resolution AFM image (Figure 5B). Previously, it has been reported that lecanemab preferentially binds to protofibrillar aggregates compared to oligomers or mature fibrils^51, 52^. The height profiles extracted along the Lecanemab-treated Aβ-42 fibrils are shown in the corresponding color-coded line profiles (Fig. 5C). In particular, the height profiles extracted along the fibril backbone (blue plot, Fig. 5D) and the underlying surface (orange plot, Fig. 5D) confirm that lecanemab does bind to the protofibrillar aggregates. The AFM-IR measurement of lecanemab-treated Aβ-42 protofibril (Figure 5E) showed similar signal intensities in the IR maps for parallel β-sheets at 1636 cm^-1^ (Figure 5F) and α-helices at 1668 cm^-1^ (Figure 5G), and no signal for the anti-parallel β-sheets at 1695 cm^-1^cm^-1^, point towards a predominantly β-sheet structure. However, the peak is shifted towards higher wavenumbers than the spectrum for pure, untreated Aβ-42. The bulk FTIR spectra also point toward fewer β-sheets after 24 hours of aggregation (Supporting Information, Figure S10). In contrast to aducanumab, the FTIR spectrum at 0.5 h had a slight shoulder at 1630 cm^-1^, and the ThT assay confirmed that lecanemab treatment of Aβ-42 peptides only led to a minor decrease in aggregation rate, which is in agreement with previous findings^21^, thus validating the results from a localized nanospectroscopy approach. To demonstrate the reproducibility of the AFM-IR measurements, we have provided additional datasets in the supporting information for untreated Aβ-42 (Fig. S11) and aducanumab (Fig. S12), and Lecanemab treated Aβ-42 protein aggregates (Fig. S13)

**Figure 5:**
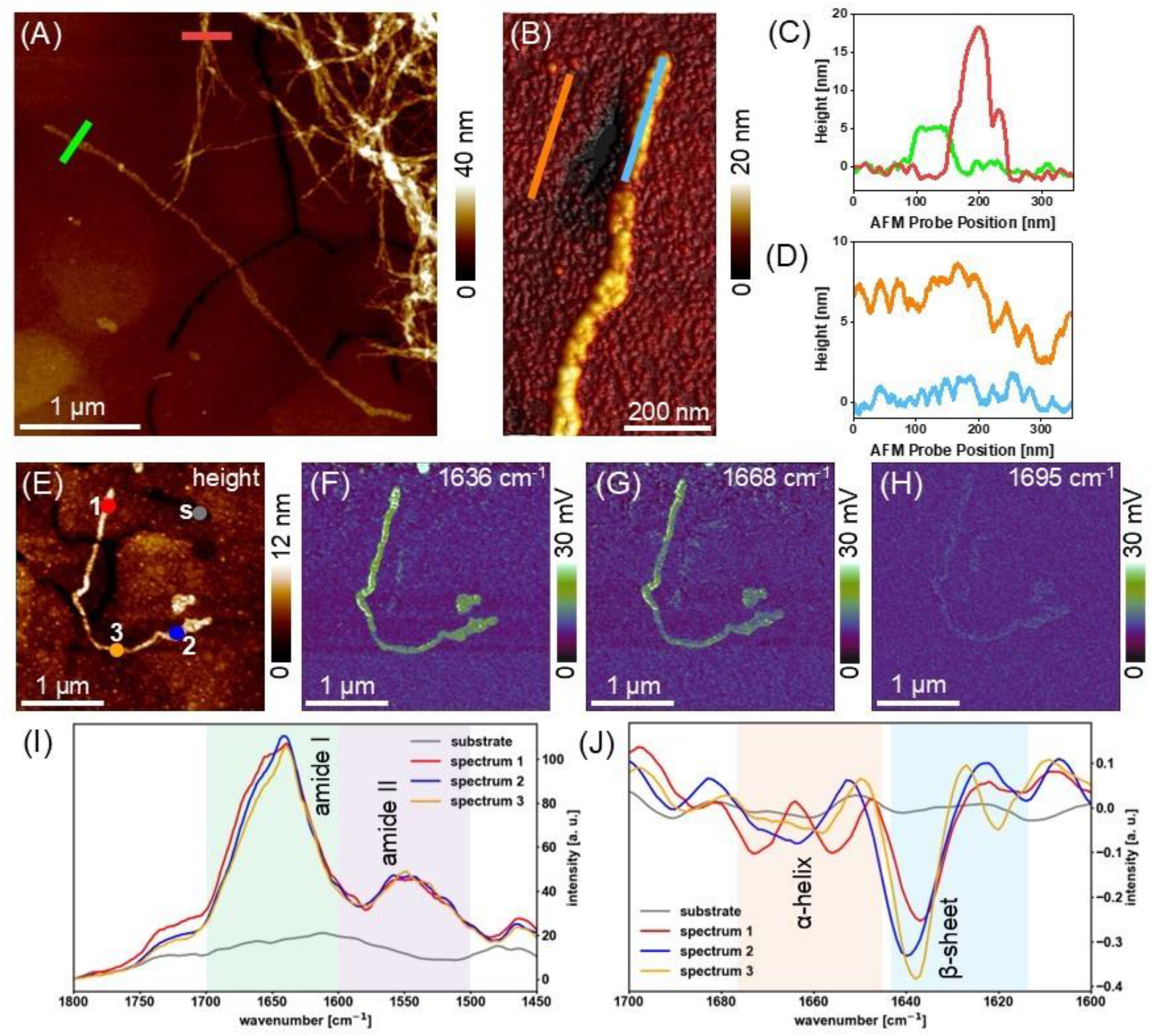
Nanospectroscopic analysis of Aβ-42 protein aggregates treated with lecanemab using tapping mode AFM and contact mode AFM-IR. (A) Tapping mode AFM height image of lecanemab treated Aβ-42 incubated for 24 h deposited on a gold substrate showing different morphologies, colored lines indicate the locations of the height profiles in (C). (B) Tapping mode AFM 3D height image of a fibril with adsorbed lecanemab, colored lines indicate the locations of the height profiles in (D). (C) Height profiles over a lecanemab decorated fibril (green) a fibril bundle (red). (D) Height profiles along the lecanemab decorated fibril (blue) and gold substrate (orange). (E) Contact mode AFM-IR height image of lecanemab treated Aβ-42 incubated for 24 h deposited on a gold substrate. (F-H) Corresponding IR maps at 1636 cm^-1^, 1668 cm^-1^, and 1695 cm^-1^, wavenumbers within the amide I region showing the contribution from parallel β-sheet, α-helix, and anti-parallel β-sheet secondary structure, respectively. (I) Representative local AFM-IR spectra for the substrate (marked “s” in the height image) and three fibrils (marked “1-3”) showing the amide I and II regions (1450-1800 cm). (J) Second derivatives of the spectra showing the amide I region, the minima correspond to the contribution from α-helix and β-sheet secondary structure.

## Conclusion

Clarifying the molecular steps involved in the aggregation of proteins implicated in neurodegenerative diseases in the presence of anti-amyloid antibodies is an area of active research interest. In this paper, we demonstrate using a combined nanoscale imaging and chemical spectroscopic approach (nanospectroscopy) to directly visualize the effects of two anti-amyloid antibodies on a wide spectrum of Aβ-42 protein aggregates. The experimental studies do not require external fluorophore labels for imaging and were conducted under standard laboratory conditions with single-particle level specificity. The main observations from this nanospectroscopy study on the differential effects of aducanumab and lecanemab on Aβ-42 protein aggregates are as follows: (1) Incubating Aβ-42 peptides together with aducanumab results in a significant reduction of free oligomer population (primary aggregation pathway). (2) aducanumab-treated Aβ-42 single fibrils tend to bundle into larger diameter fibrils, confirmed by AFM height maps. (3) Incubating Aβ-42 peptides together with lecanemab did not lead to quenching of oligomer generation. (4) Lecanemab displayed a higher affinity for Aβ-42 protofibrils, evidenced by the morphologically distinct fully-coated protofibrillar structures. (5) In contrast to known levodopa-induced disassembly of Aβ-42 fibrils to form off-pathway oligomers^53, 54^, both aducanumab-treated and lecanemab-treated Aβ-42 fibrils and protofibrils did not disassociate into smaller fragments. We anticipate that the insights gained from this study on anti-amyloid antibodies interacting with specific protein targets could serve as useful guidelines in developing immunotherapies that are designed to be effective even in preclinical or asymptomatic stages of AD. Furthermore, the nanospectroscopy approach demonstrated in this work is not limited to only probing synthetically prepared proteins but can also be extended to characterise recombinant protein aggregates such as amyloid and tau isoforms in the CSF of AD patients and compared with individuals who have received the AD drugs in clinical trials. Knowledge gained from such studies will not only help provide clues on disease progression but also verify in a label-free manner the effectiveness of the prescribed treatment in individuals who have received the AD drugs in clinical trials.

## Associated Content

### Materials and Methods

#### Amyloid beta solution synthesis

Synthetic Aβ-42 was purchased from Abcam and processed as described in a previous study^14^. One vial containing 1 mg of Aβ-42was dissolved in 1 mL 10% (V/V) ammonium hydroxide, divided into 25 μg aliquots, and freeze-dried. The aliquots were stored at ™20°C until further use. A stock solution was prepared by dissolving the freeze-dried pellet in 50 μL 60 mM NaOH, the exact concentration was determined by UV spectroscopy (Implen NanoPhotometer) using the extinction coefficient of 1490 M^-1^cm^-114^. The aggregation was started by diluting the stock solution with PBS to reach a monomer concentration of 5 μM and incubating the sample at 37°C. To observe the binding of antibodies to the Aβ-42 aggregates, 100 μL of the 5 μM peptide solution was mixed with 100 μL aducanumab (Proteogenix) or lecanemab (ichorbio) solution at a concentration of 20 μg/mL, resulting in a final Aβ-42 concentration of 2.5 μM and 10 μg/mL antibody as reported in a previous study^21^.

#### Atomic force microscopy measurements

After aggregation from 4 to 96 hrs, 5 μL of the Aβ-42 untreated and antibody-treated samples were deposited on a gold substrate (purchased from Phasis Inc) and air-dried overnight at room temperature. The excess salt adsorbates were rinsed off after drying using 1 mL water and the residual water was gently blown off with a small hand pump. Tapping mode AFM was performed on the treated and untreated Aβ-42 samples using a Bruker Dimension Icon instrument with NuNano SCOUT 70 HAR RAu probes (70 kHz resonance frequency, spring constant 2 Nm^-1^, reflective gold coating, nominal radius 5 nm). Images were recorded at typical scan rates of 0.5 Hz and 1024×1024 lines. The raw data was processed (1st order flattened and low pass filtered) and analyzed using the Bruker NanoScope Analysis software.

#### Infrared nanospectroscopy (AFM-IR)

The same treated and untreated Aβ-42 samples on gold substrates (enhancing the IR signal^56^) were imaged with an Anasys NanoIR 2 system equipped with Bruker PR-EX-TnIR-A tapping mode probes (75 kHz resonance frequency, spring constant 1-7 Nm^-1^, gold coating on the cantilever and tip with a nominal radius of 20 nm) and Bruker PR-EX-CnIR-B contact mode probes (13 kHz resonance frequency, spring constant 0.1-0.4 Nm^-1^, gold coating on the cantilever and tip with a nominal radius of 20 nm). Contact mode measurements were performed in resonance-enhanced mode, tuning the laser pulses at a repetition rate matching the first contact resoncance frequency of the cantilever. The AFM maps were recorded at a scan rate of 0.5 Hz and at a resolution of 512×512 lines. The raw data was flattened with the Gwyddion software (0^th^ to 2^nd^ order flattening). A quantum cascade laser was used as the IR source and 5 spectra at a resolution of 1 cm^-1^ were averaged. To reduce water bands in the spectra, the relative humidity was kept below 5% during the measurements using a constant flux of N_2_. The recorded spectra were treated using the Anasys Analysis Studio to eliminate step discontinuity filter to compensate for offsets at the laser chip transitions. The raw spectra were further processed using Python. Spikes deviating more than 30% from the mean of the 7 neighboring data points were eliminated and the spectra were smoothed with a 2^nd^ order Savitzky-Golay filter with a window of 25 points. The 2^nd^ derivatives of the amide I region of the final smoothed spectra were computed to better visualize the different secondary structure contributions.

#### KPFM measurements

Local measurements of the surface-potential distribution were performed on the aducanumab-treated Aβ-42 and untreated Aβ-42 sample using scanning Kelvin probe force microscopy (SKPFM). To this end, a Bruker ICON 3 atomic force microscope (AFM) was used in Peak Force Scanning Kelvin Probe Force Microscopy (PF-SKPFM) mode with a conductive Bruker PFQNE-AL probe (300 kHz resonance frequency, nominal spring constant 0.8 Nm^-1^, highly doped silicon tip with a nominal radius of 5 nm, SiN_3_ cantilever with aluminum-based conductive coating on the back side) at a scanning rate of 0.3 Hz. The probe was scanned in a dual pass over the sample surface, whereby line-by-line height profile and surface potential profile are sequentially acquired. Hereby, surface potential images were obtained in lift mode at a constant lift scan height (tip-sample distance) of 10 nm. Measurements were performed at RT. If not stated otherwise, a tip bias of 1 V was applied during the measurement, which proved yielding the most favorable signal-to-noise ratio without altering the sample. Images were acquired on single fibers to exclude signal contribution and possible interference by overlapping or crossing fibers. If not stated otherwise, KPFM measurements were analyzed and reported as measured, to ensure internal consistency and avoid potential uncertainties introduced by external referencing. The interpretation focuses on relative variations within a sample, providing a robust comparison of potential distribution between sample conditions.

Prior to imaging, samples deposited on Au-film/mica substrates were mounted on conductive stainless-steel discs and their Au surface electrically contacted with silver paste.

Measurements were performed on individual fibers to avoid artefacts that may arise for crossing or overlapping Aβ-42 nanofibrils [G. Lee, W. Lee, H. Lee, S. W. Lee, D. S. Yoon, K. Eom, and T. Kwon, Appl. Phys. Lett., 101, 043703 (2012)].

#### ATR-FTIR spectroscopy

Attenuated total reflectance Fourier transform infrared spectroscopy (ATR-FTIR) was conducted using a Bruker Tensor-27 spectrometer. After obtaining background measurements, the two times 5μL of the amyloid suspensions were dried on the ATR diamond using a stream of air. The spectra were recorded at a resolution of 4 cm^-1^ and with 32 co-averages.

#### Zeta potential measurements

The zeta potential of untreated and treated Aβ-42 samples was measured with a Malvern Zetasizer equipped with a folded capillary cell (Malvern DTS1070). Due to the high ionic strength and resulting conductivity in PBS, the measurements were conducted with the recommended diffusion barrier method to prevent corrosion on the electrodes of the cell^57^. Measurements were attempted in triplicate for each condition; however, only the first run of each measurement provided high-quality data. Subsequent measurements showed low count rates and degraded phase signal, likely due to fibril migration toward the electrodes, despite using the diffusion barrier method. Therefore, only the first runs from each of two independent triplicate measurements was included in the final analysis. These values are reported as the average of two independent first-run measurements, without standard deviation due to the limited number of replicates.

#### Thioflavin T fluorescence aggregation kinetics assay

Identical concentrations (2.5 μM Aβ-42 and 10 μg/mL aducanumab or lecanemab) for the AFM-based experiments were used to study the effect of the antibodies on the aggregation kinetics. The cross-β-sheet specific fluorescent dye Thioflavin T was added at a final concentration of 5 μM. The ThT fluorescence was measured using a BMG Labtech CLARIOstar Plus plate reader, Greiner 96-well plates with F-bottom sealed with foil to prevent evaporation. The plate was measured every 60 s from the bottom, using an excitation wavelength of 440 nm and detecting emission at 482 nm. The measurements were repeated twice, each time performed on triplicates. The average of the triplicate and the standard deviation were calculated for the Aβ-42 with and without antibody treatment and the pure antibodies at the same concentration. A blank of pure ThT was subtracted from each sample.

## Author Contribution

N.K. performed the AFM and AFM-IR measurements, and M.C. conducted the KPFM measurements. P.N.N designed the study and was involved in the analysis of AFM, AFM-IR, and KPFM data. N.K. and P.N.N. wrote the manuscript. All authors discussed the results and commented on the manuscript.

## Notes

The authors declare no competing financial interests

## Acknowledgments

Funding: P.N.N and N.K. thank the Lazarus-Stiftung Foundation, Sanare, and Theodor Naegeli-Stiftung for their financial support. M.C. acknowledges support from the Swiss National Science Foundation (SNSF) via an Ambizione Fellowship (Grant No. PZ00P2_216383). N.K thanks Prof. Dr. Nancy Burnham for valuable discussions, the Particle Laboratory at Eawag for granting access to Malvern Zetasizer for zeta potential measurements, and the Functional Polymer Laboratory at Empa for access to the FTIR spectrometer.

