## Supplementary Information for "Anti-amyloid antibody effects on Aβ-42 protein aggregates profiled using nanospectroscopy"

**Contents**

Figure S1: Quantifying Aβ-42 oligomer particle size from AFM height maps
Figure S2: Mapping surface potential distribution of amyloid Aβ-42 aggregates with and without aducanumab treatment.
Figure S3: Assessment of Kelvin Probe Force Microscopy (KPFM) acquisition parameters.
Figure S4: Surface potential variations along a single adsorbed amyloid Aβ-42 fiber with and without aducanumab treatment.
Figure S5: High-resolution mapping of aducanumab adsorbed on Aβ-42 fibril surface
Figure S6: Raw, de-spiked and smoothed (2nd order, 25 point Savitzky-Golay) spectra corresponding to the spectra shown in Figure 3I.
Figure S7: Fourier-transform infrared spectra of Aβ-42 samples with and without aducanumab treatment after 0.5 h and 24 h of incubation.
Figure S8: Raw, de-spiked, and smoothed (2nd order, 25 point Savitzky-Golay) spectra corresponding to the spectra shown in Figure 4I.
Figure S9: Thioflavin T (ThT) fluorescence as a function of time for pure Aβ-42 samples and samples incubated with aducanumab (red) and Lecanemab.
Figure S10: Fourier-transform infrared spectra of Aβ-42 samples with and without lecanemab treatment after 0.5 h and 24 h of incubation.
Figure S11: Additional nanospectroscopic analysis of Aβ-42 protein aggregates using contact mode AFM-IR.
Figure S12: Additional nanospectroscopic analysis of Aβ-42 protein aggregates treated with aducanumab using contact mode AFM-IR.
Figure S13: Additional nanospectroscopic analysis of Aβ-42 protein aggregates treated with lecanemab using contact mode AFM-IR.

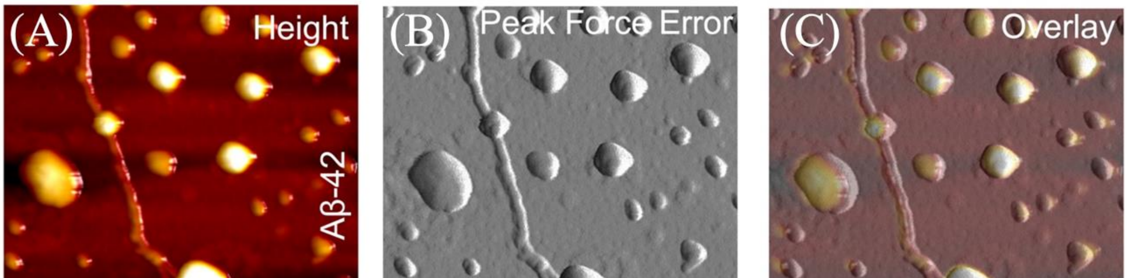

**Figure S1: Quantifying Aβ-42 oligomer particle size from AFM height maps.** (A) AFM height map of purified Aβ-42 protein aggregates and simultaneously acquired peak force error (B), showing the presence of both spherical and fibrillar particles. (C) Overlay of AFM height and peak force error from panels A and B. (D-I) A series of height profiles extracted from a single spherical particle, which could potentially represent Aβ-42 oligomers, to obtain the spherical particle heights.

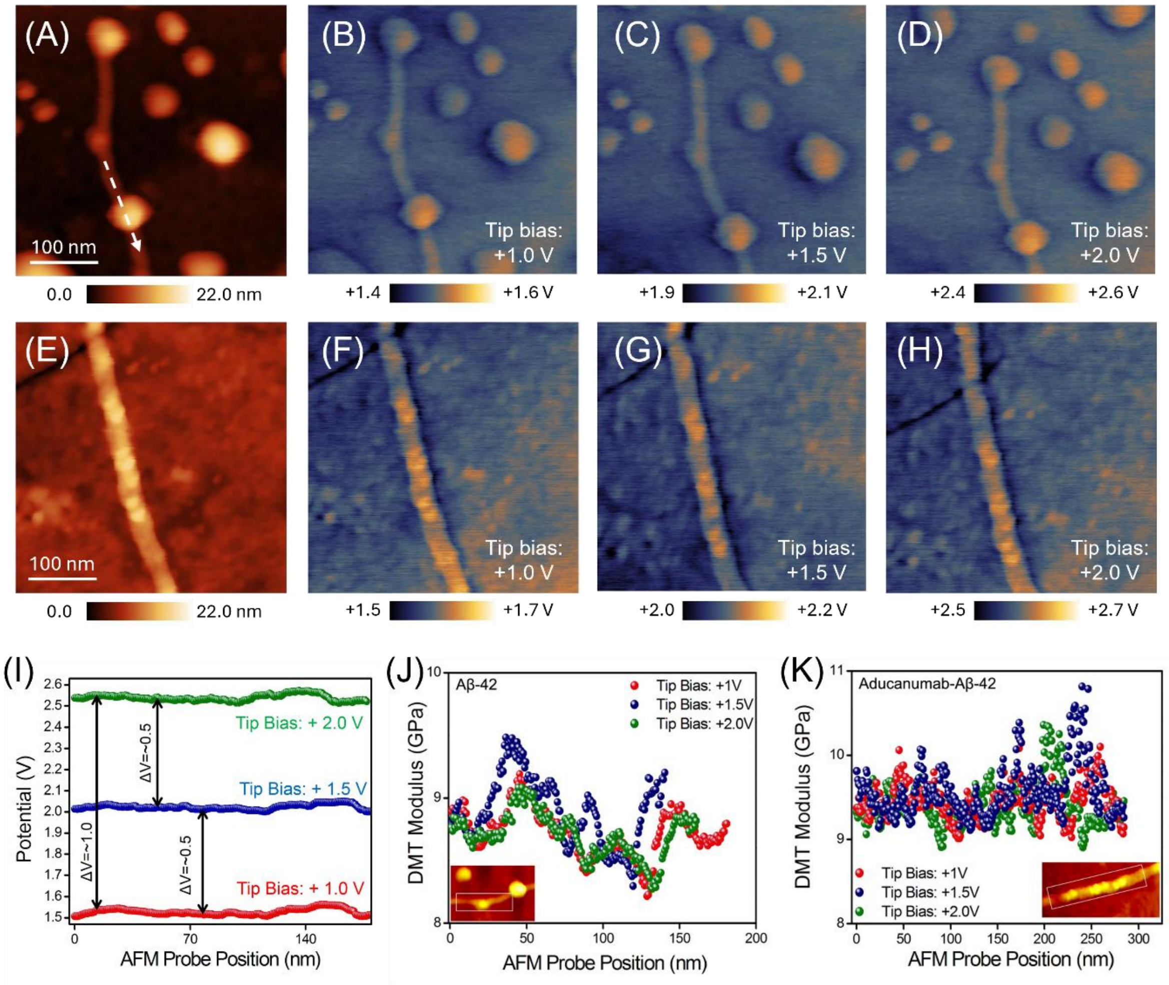


**Figure S2: Mapping surface potential distribution of amyloid Aβ-42 aggregates with and without aducanumab treatment.** (A) Height map of purified Aβ-42 protein aggregates showing both fibrillar and spherical particles. (B-D) KPFM maps were acquired simultaneously (line-by-line dual pass) as the height map shown in panel A at +1.0 V, +1.5 V, or +2.0 V tip bias, respectively. (E) Height map of aducanumab-treated Aβ-42 protein aggregates showing mostly fibrillar aggregates of larger diameter. (F-H) A series of KPFM maps acquired simultaneously (line-by-line dual pass) as the height map shown in panel E, at +1.0 V, +1.5 V, or +2.0 V tip bias, respectively. (I) extracted line profile across a purified Aβ-42 protein aggregate fiber with spherical particle (location marked with a white line in (A). The measured surface potential shifts near-linearly with the applied tip bias, while the relative potential differences across the amyloid fiber (~0.5 V intra-fibril variation) remain consistent at each bias. This indicates minimal tip-sample interaction artifacts and confirms the reliability of the potential measurements. (J+K) DMT modulus as a function of AFM probe position along the region marked in the white box shown in the lower left inset AFM image of (J) purified and (K) aducanumab-treated Aβ-42 protein aggregates for different tip biases. The mechanical response is revealed to be largely independent from the applied electrical field, with only a slight tendency for bias-induced softening for the purified fiber (J). The overall modulus appears higher for the aducanumab-treated condition (K), suggesting structural stabilization from aducanumab binding.

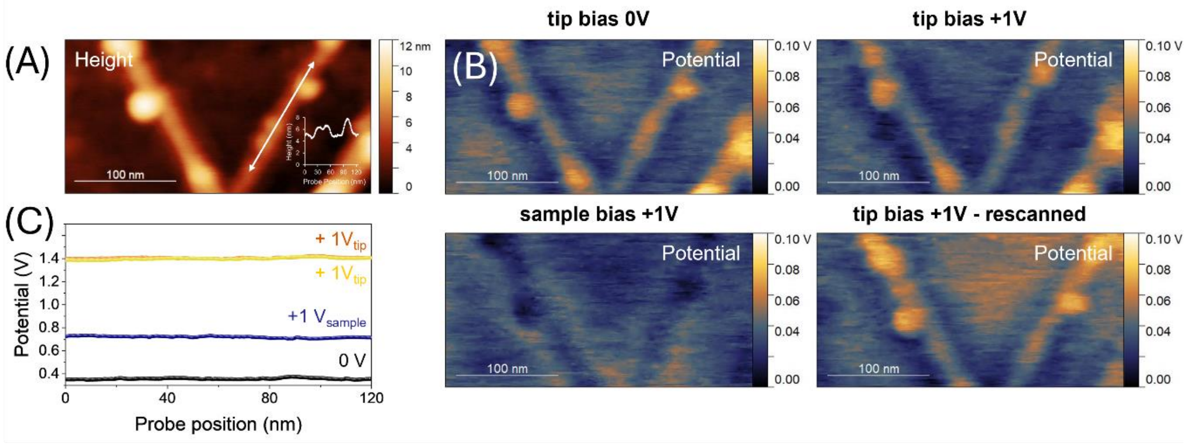


**Figure S3: Assessment of Kelvin Probe Force Microscopy (KPFM) acquisition parameters.** (A) Height map of purified Aβ-42 protein aggregates showing both fibrillar and spherical particles. (B) Surface potential distribution images at varying acquisition conditions: Applying +1V tip bias stabilizes the signal without altering the intrinsic charge distribution of the sample and yields a favorable signal-to-noise ratio, thus revealing the highest level of detectable potential variation across a single fiber. In contrast, an applied sample bias of +1V considerably worsens the signal and detailed information of potential variation is lost, indicating provocation of charge redistribution or charge injection, effectively screening local potential variations. Rescanning of the same area at a tip bias of +1V, following +1V sample bias, shows that the charge distribution in the exposed field of view remained largely unaffected, with a small loss in resolution, which is ascribed to charge modification by the preceding sample bias. All potential images were normalized for comparability between the different scales. (C) Surface potential line profiles: potential variation across a line for the different potential biases applied (location of line marked with a white arrow in the height image) displays the effectively measured potentials. Applying a tip bias resulted in a close to equivalent shift in the measured potential (electrostatic potential responds nearly linearly to applied voltage), indicating that the electrostatic interaction is preserved. In contrast, applying a bias to the sample resulted in a significantly damped measured potential, supporting the assumption of charge screening, and capacitive or charge redistribution effects of the thin molecular layer.

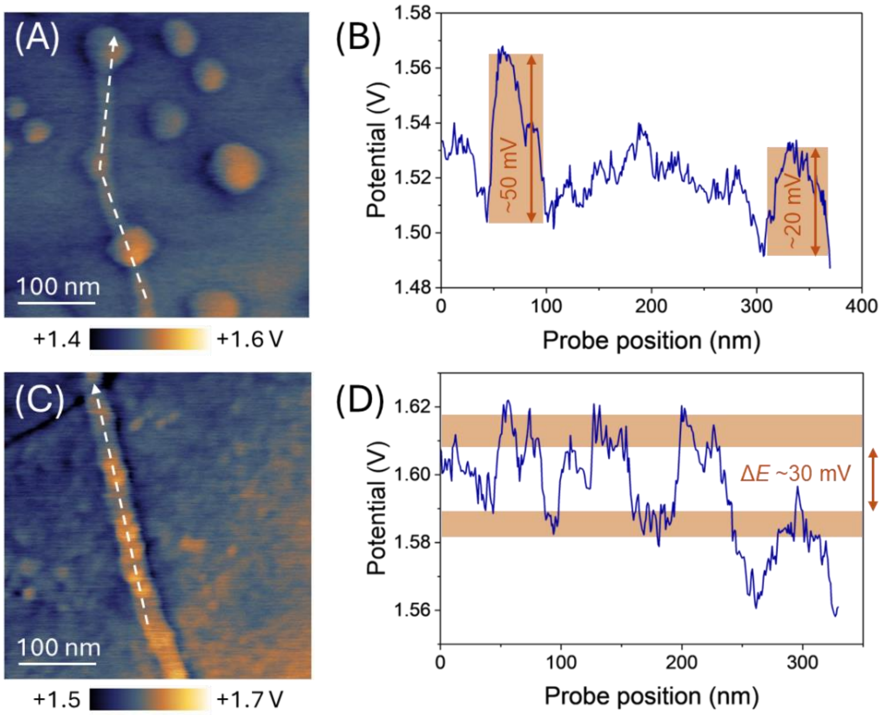


**Figure S4: Surface potential variations along a single adsorbed amyloid Aβ-42 fibril with and without aducanumab treatment.** (A-C) Surface potential map at +1.0 V tip bias without (A) and with (C) aducanumab treatment, with the dashed white lines indicating the scan paths used for potential profiling. (B+D) Corresponding potential profiles extracted along the fibrils in (A) and (C), respectively, showing moderate local variations in surface potential, on the order of tens of millivolts. The similarity in magnitude across both conditions suggests consistent electrostatic behavior of the fibrils.

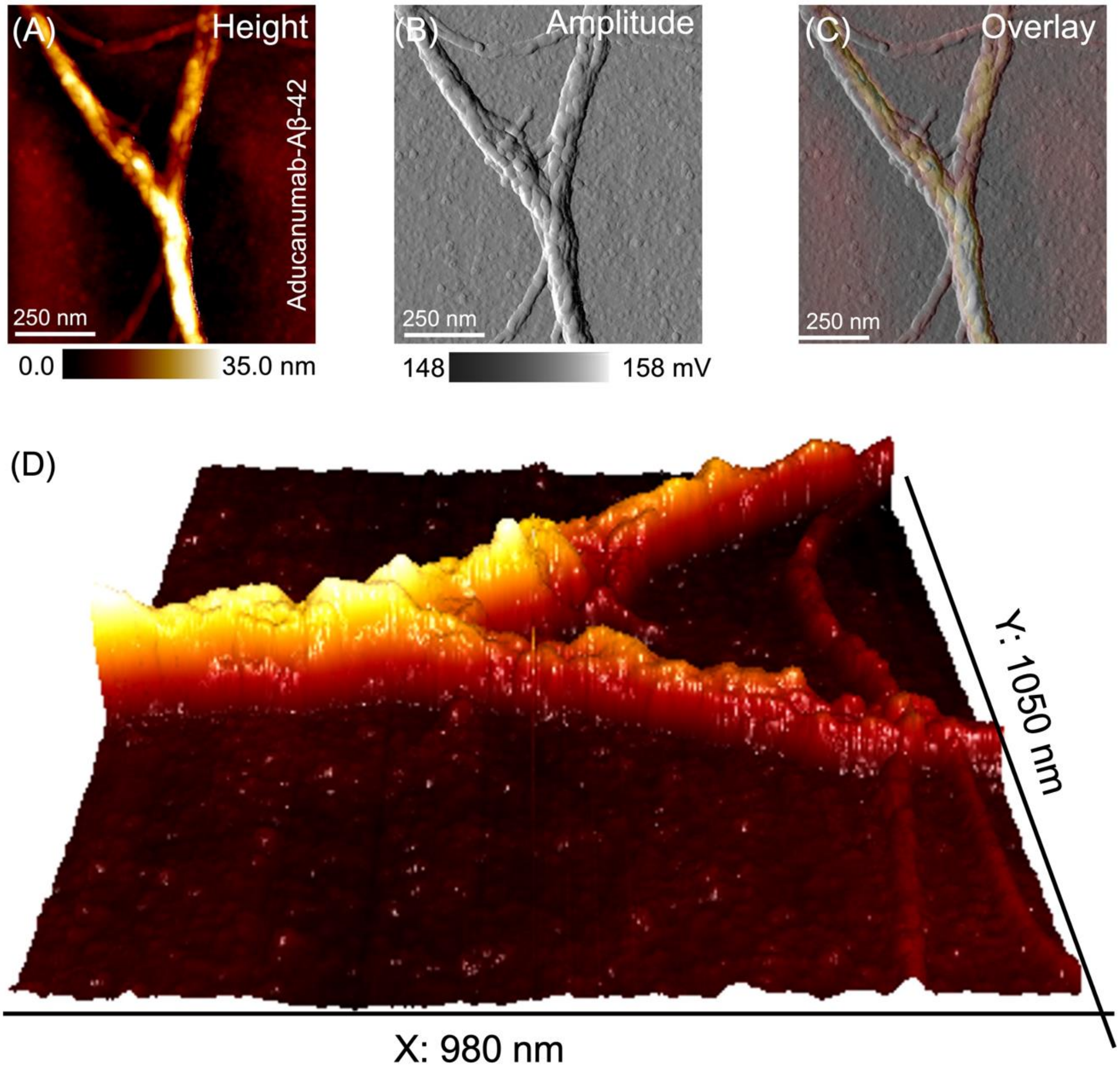


**Figure S5: High-resolution mapping of aducanumab adsorbed on Aβ-42 fibril surface.** (A) AFM height map of aducanumab-treated Aβ-42 protein aggregates showing mostly fibrils assembled in a bundled manner in contrast to single fibrils. (B) Simultaneously acquired amplitude image showing the aducanumab molecules adsorbed along the length of the Aβ-42 fibril bundle surface. (C) Overlay image of height and amplitude shown in panels A and B. (D) Three-dimensionally represented height map of the Aβ-42 fibrils showing clearly the bundle morphology compared to single fibrils also resolved within the same AFM frame.

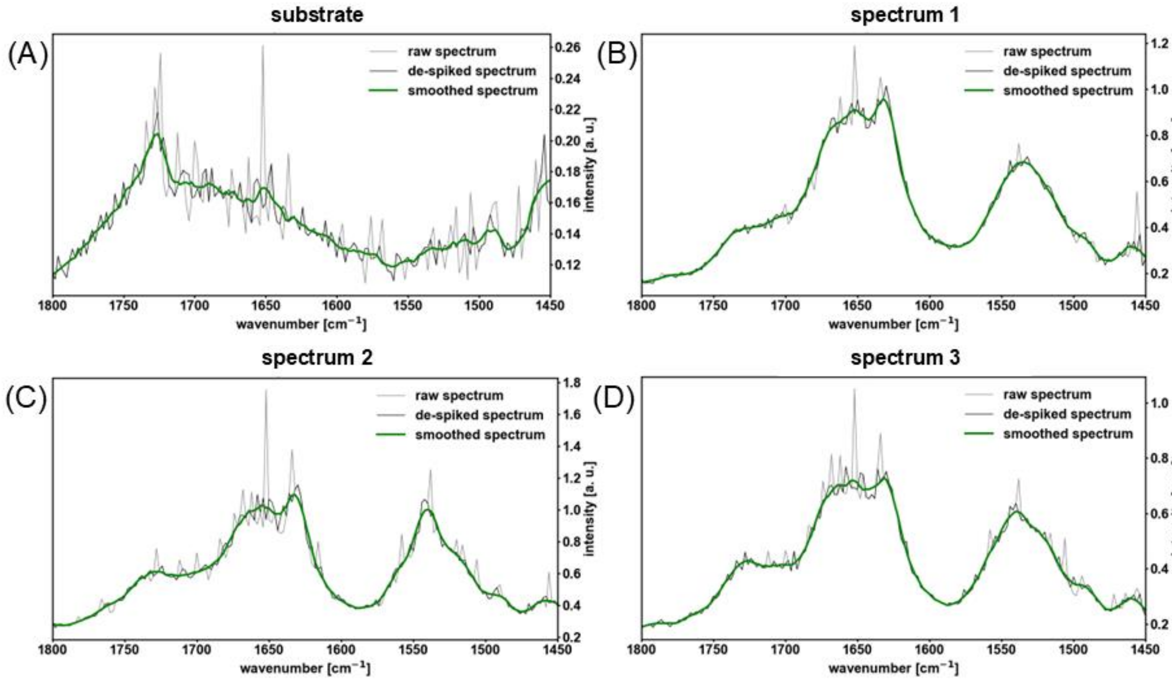


**Figure S6**: (A-D) Raw, de-spiked and smoothed (2nd order, 25 point Savitzky-Golay) spectra corresponding to the spectra shown in Figure 3I.

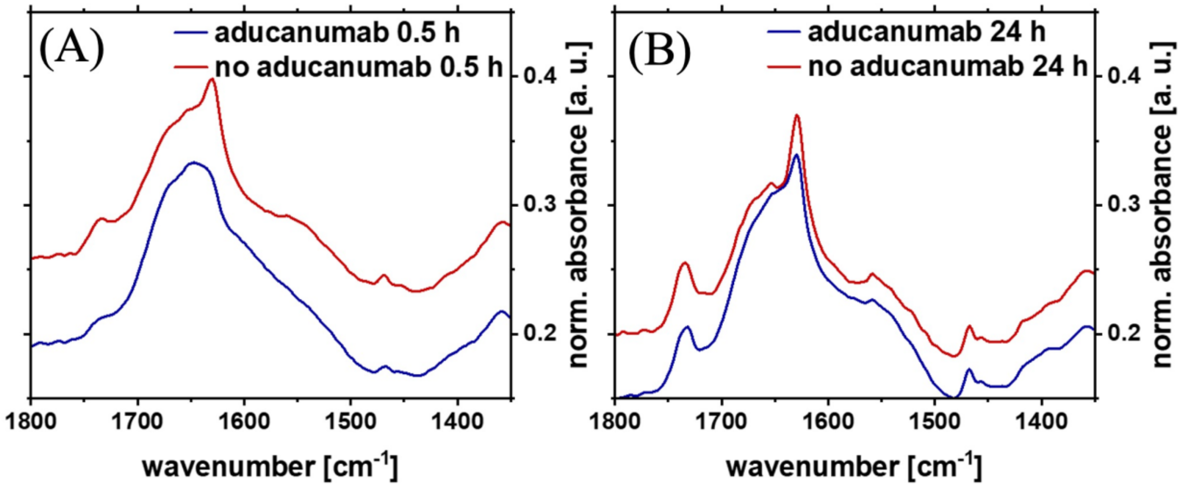


**Figure S7**: Fourier-transform infrared spectra of Aβ-42 samples with and without aducanumab treatment after 0.5 h and 24 h of incubation.

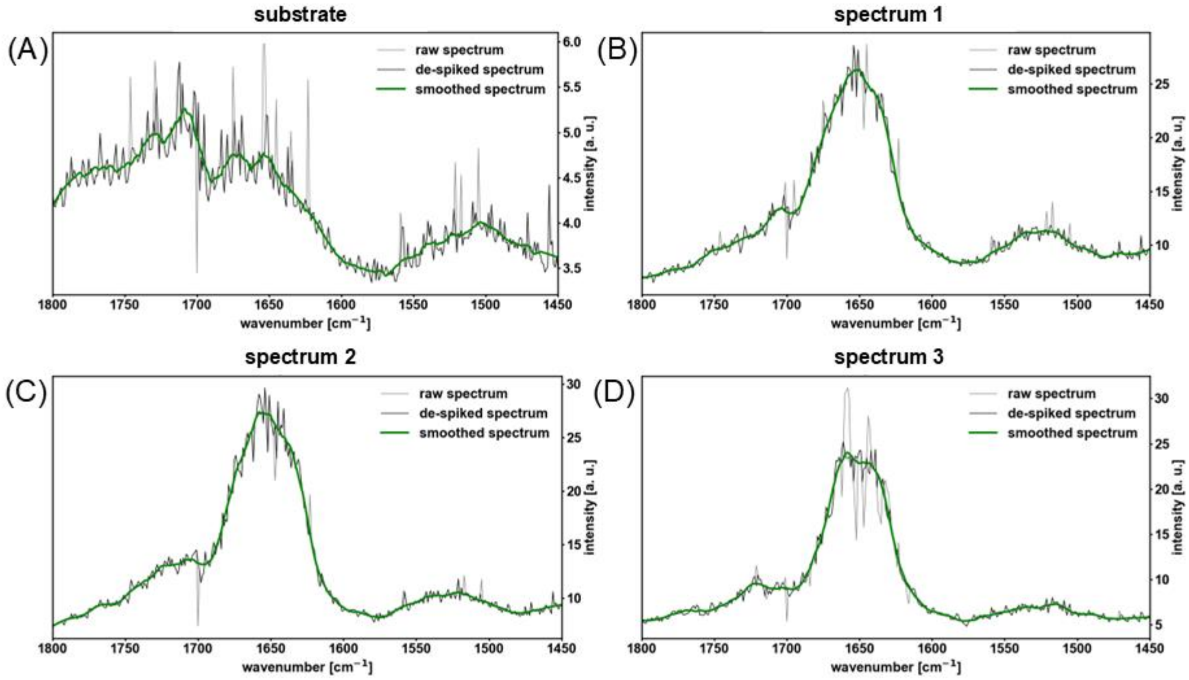


**Figure S8:** (A-D) Raw, de-spiked, and smoothed (2nd order, 25 point Savitzky-Golay) spectra corresponding to the spectra shown in Figure 4I.

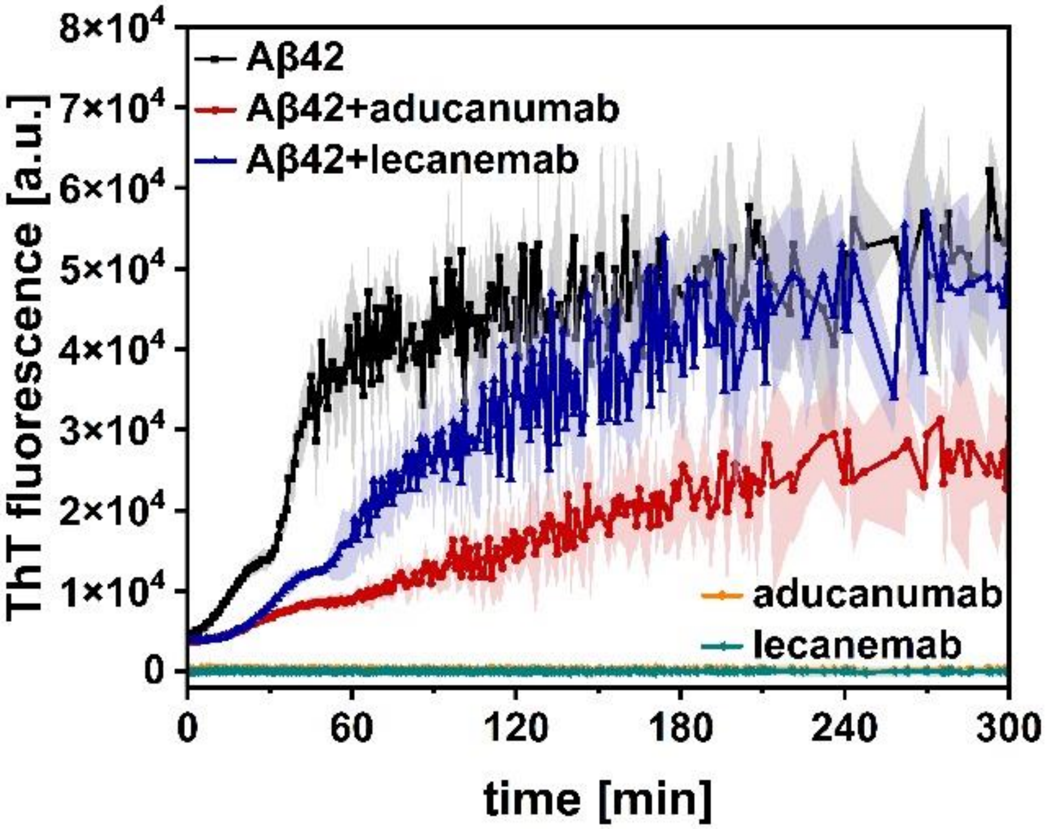


**Figure S9**: Thioflavin T (ThT) fluorescence as a function of time for pure Aβ-42 samples (black) and samples incubated with aducanumab (red) and Lecanemab (blue). The shown data points of averaged triplicates come with a standard deviation in a lighter color shadow. The antibody controls did not show any increase in fluorescence over time. A blank of pure ThT at equal concentration of the samples and controls was subtracted.

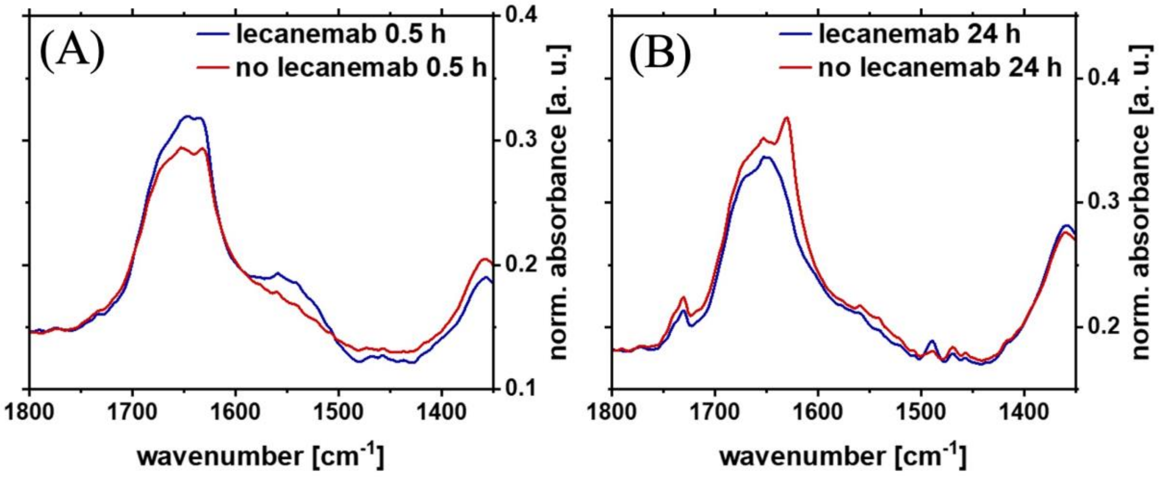


**Figure S10:** Fourier-transform infrared spectra of Aβ-42 samples with and without lecanemab treatment after 0.5 h and 24 h of incubation.

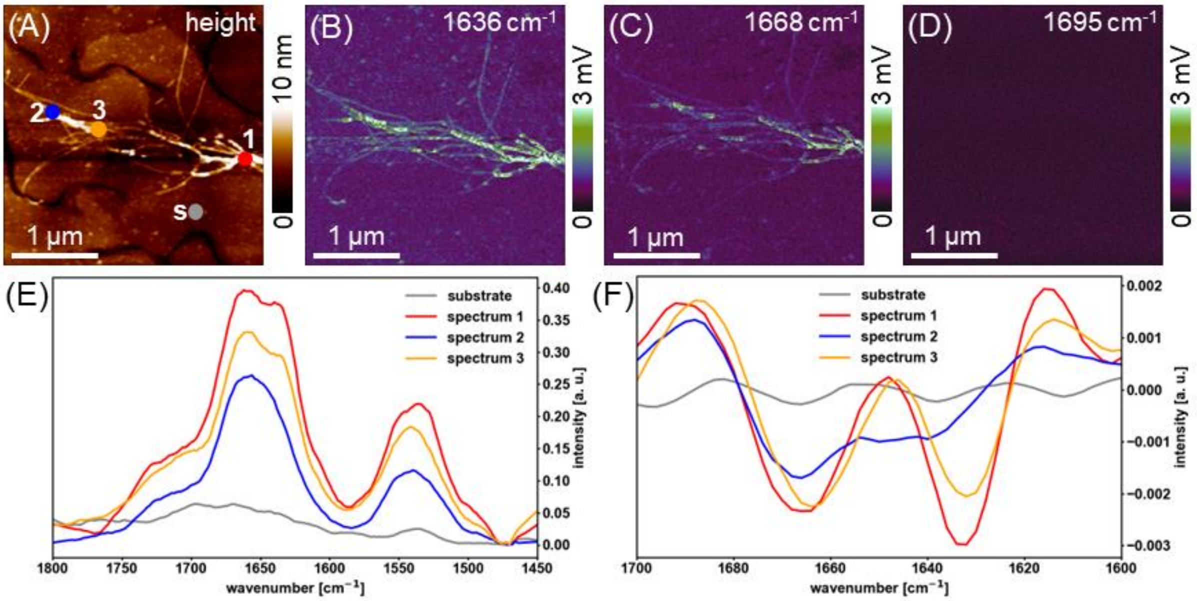


**Figure S11: Additional nanospectroscopic analysis of Aβ-42 protein aggregates using contact mode AFM-IR**. (A) Contact mode AFM-IR height image of untreated Aβ-42 incubated for 24 h deposited on a gold substrate. (B-D) Corresponding IR maps at 1636 $cm^{-1}$, 1668 $cm^{-1}$ and 1695 $cm^{-1}$, wavenumbers within the amide I region showing the contribution from parallel β-sheet, α-helix and anti-parallel β-sheet secondary structure, respectively. (E) Representative local AFM-IR spectra for the substrate (marked "s" in the height image) and three fibrils (marked "1-3") showing the amide I and II regions (1450-1800 cm). (F) Second derivatives of the spectra showing the amide I region, the minima correspond to the contribution from α-helix and β-sheet secondary structure.

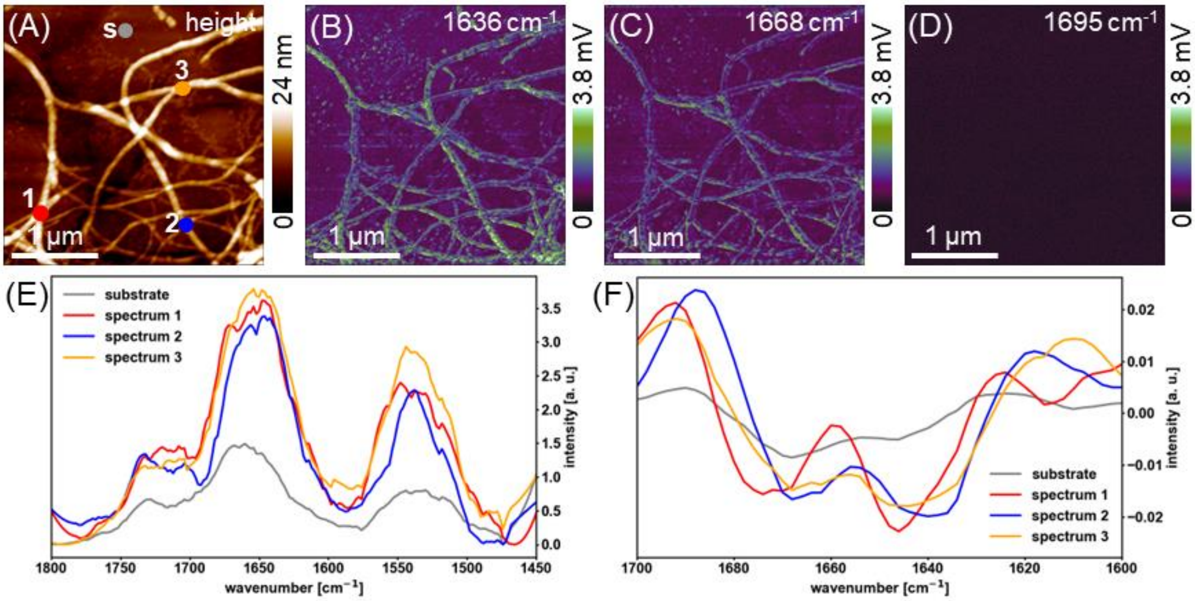


**Figure S12: Additional nanospectroscopic analysis of Aβ-42 protein aggregates treated with aducanumab using contact mode AFM-IR**. (A) Contact mode AFM-IR height image of Aβ-42 treated with aducanumab incubated for 24 h deposited on a gold substrate. (B-D) Corresponding IR maps at 1636 $cm^{-1}$, 1668 $cm^{-1,}$ and 1695 $cm^{-1}$, wavenumbers within the amide I region showing the contribution from parallel β-sheet, α-helix, and anti-parallel β-sheet secondary structure, respectively. (E) Representative local AFM-IR spectra for the substrate (marked "s" in the height image) and three fibrils (marked "1-3") showing the amide I and II regions (1450-1800 cm). (F) Second derivatives of the spectra showing the amide I region, the minima correspond to the contribution from α-helix and β-sheet secondary structure.

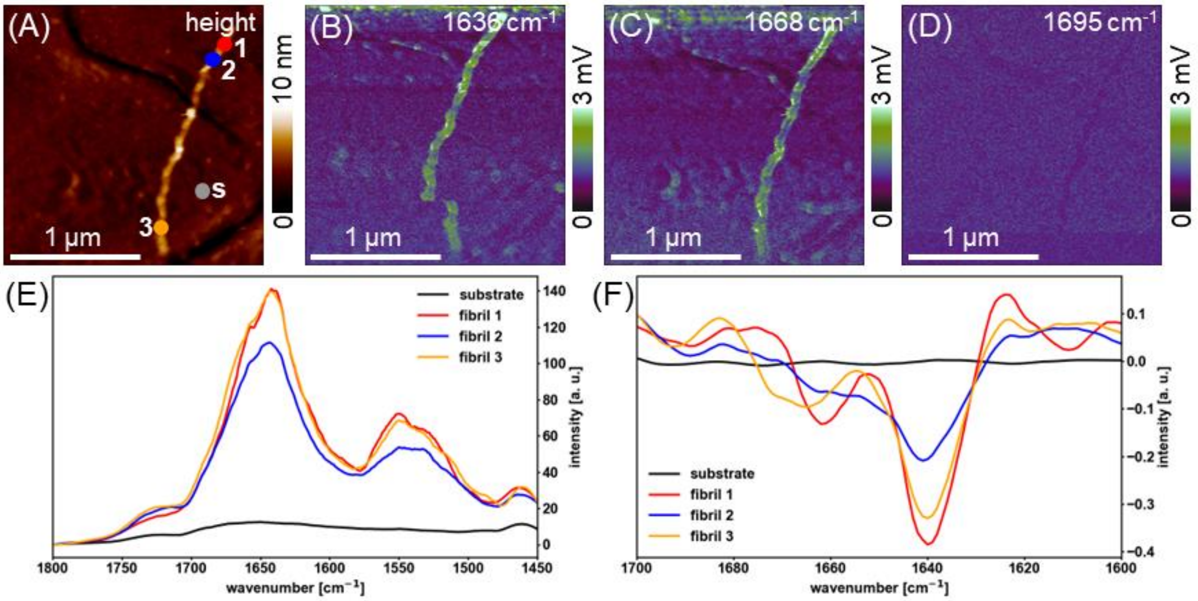


**Figure S13: Additional nanospectroscopic analysis of Aβ-42 protein aggregates treated with lecanemab using contact mode AFM-IR**. (A) Contact mode AFM-IR height image of Aβ-42 treated with lecanemab incubated for 24 h deposited on a gold substrate. (B-D) Corresponding IR maps at 1636 $cm^{-1}$, 1668 $cm^{-1}$ and 1695 $cm^{-1}$, wavenumbers within the amide I region showing the contribution from parallel β-sheet, α-helix and anti-parallel β-sheet secondary structure, respectively. (E) Representative local AFM-IR spectra for the substrate (marked "s" in the height image) and three fibrils (marked "1-3") showing the amide I and II regions (1450-1800 cm). (F) Second derivatives of the spectra showing the amide I region, the minima correspond to the contribution from α-helix and β-sheet secondary structure.
